# *Trichomonas vaginalis* pseudocysts are a quiescent cell stage characterized by a differentially regulated transcriptome and intact membranes

**DOI:** 10.64898/2026.06.14.730637

**Authors:** Mari Shiratori, Francisco Callejas-Hernández, Jordan C. Orosco, Steven A. Sullivan, Carlos Carmona Fontaine, Jane M. Carlton

**Affiliations:** Center for Genomics and Systems Biology, Department of Biology, New York University, New York City, NY, United States of America; Department of Molecular Microbiology and Immunology, Bloomberg School of Public Health, Johns Hopkins University, Baltimore, MD 21205, United States of America; Department of Biomedical Engineering, Whiting School of Engineering, Johns Hopkins University, Baltimore, MD 21218 USA

**Author notes:** Corresponding author: (JMC).

## Abstract

*Trichomonas vaginalis* is the causative agent of trichomoniasis, the most common non-viral sexually transmitted infection (STI). Despite its prevalence, low levels of public knowledge and research funding and the absence of *T. vaginalis* screening or control programs have led to its categorization as a “neglected” STI. Unlike other STIs in the USA, prevalence increases with age, peaking among individuals in their 40s. Both a motile trophozoite stage and a non-motile pseudocyst state have been described for the parasite, although it is debated whether the latter is a quiescent stage or a degenerate form on its way to cell death. Here we characterize the *T. vaginalis* pseudocyst by flow cytometry, membrane integrity assays, transcriptomics, and reversion studies. Pseudocysts were induced by culturing trophozoites in acidic media or in iron depleted media, with a variety of resulting survival rates. Flow cytometry studies showed that pseudocysts have intact cell membranes and express phosphatidylserine on their cell surface. Fluorescence-activated cell sorting studies also identified distinct sub-populations of parasites, revealing the importance of using pure live pseudocyst cultures in reversion studies. Pseudocysts were transcriptionally active for several days and had consistent subsets of genes with increased expression compared to trophozoites, although decreased transcription of genes involved in metabolism. Comparative transcriptomics of pseudocysts and trophozoites of two *T. vaginalis* strains revealed distinct cell states. Combined, our results provide evidence that the pseudocyst cell state is a stress-induced quiescent stage of *T. vaginalis* that can remain viable for days, with implications for a role in persistent infections.

**Author Summary:** The sexually transmitted parasite *Trichomonas vaginalis* has two well-known cell forms: a free-swimming, flagellated trophozoite and an amoeboid form adhered to host epithelia. A third morphology, the pseudocyst, has been described, but it is unclear whether this is a quiescent stage capable of facilitating persistent infections, or a degenerate form indicating cell death. Here we describe experiments revealing that pseudocysts have overall decreased gene expression compared to trophozoites but exhibit stage-specific gene transcription and plasma membrane integrity for days. Moreover, pseudocysts maintain externalized phosphatidylserine for multiple days. Taken together, our results suggest that pseudocysts are a viable cell stage in *T. vaginalis* distinct from cell death. Further research into pseudocysts and particularly their interactions with host immune cells is needed to reveal what role they may play in *T. vaginalis* pathogenesis and persistent or asymptomatic infections.

## Introduction

*Trichomonas vaginalis* is the causative agent of the most common nonviral sexually transmitted infection (STI) globally, trichomoniasis [1]. A notable characteristic of trichomoniasis is that, unlike other STIs, prevalence increases with age, peaking among patients in their 40s [2]. Many cases of trichomoniasis are asymptomatic [3], but when symptoms occur they include a burning sensation during urination, discharge with an unpleasant odor, and characteristic puncta on cervical epithelial tissue called “strawberry cervix” [4]. *T. vaginalis* has also been reported to cause persistent infections, with symptoms reappearing up to a year after treatment [5–7]. Persistent cases and the higher prevalence in perimenopausal and menopausal individuals raise questions about their underlying mechanisms.

*T. vaginalis* has three cell states: trophozoite, amoeboid, and pseudocyst. The trophozoite is free-swimming and pear-shaped, with four anterior flagella, a posterior axostyle, and an undulating membrane along its length. After infection, trophozoites adopt an amoeboid form that adheres to and migrates on host epithelium [8]. The third form observed in trichomonad species is immotile, smaller, and round, having resorbed its axostyle and flagella, and lacking a true cyst wall. This form was controversial a century ago, having been variously called ‘cyst’, ‘pseudocyst’, ‘degenerate individual’, or another parasite entirely [9]. The term ‘pseudocyst’ prevailed, although later groups have championed ‘endoflagellar form’ [10] to make internalized flagella its hallmark, and ‘cyst-like structure’ when chitin was found in the pseudocyst wall [11, 12]. The pseudocyst form became accepted as a trichomonad response to unfavorable environmental conditions, but whether it was a protective life-cycle stage or a degenerative, moribund cell generated debate, particularly for *T. vaginalis* [10, 13–16].

That pseudocysts occur in many trichomonad species (reviewed in [17]) argues for them being functional, but support for this view has varied by species. A function as a resistant form rather than degenerative was considered more plausible for gastrointestinal species where pseudocysts passed to feces or water could survive long enough to contaminate a new amphibian, reptile, bird, or rodent host, than for urogenital species passed sexually with minimal exposure to harsh external environments. But strong evidence for viability of the latter emerged from studies of pseudocysts of the bovine venereal disease agent *Tritrichomonas foetus* that colonizes bull foreskins and cow vaginas. Granger et al., [18] reversibly induced such pseudocysts *in vitro* by cooling and rewarming, and also observed pseudocysts in standard culture. *Tr. foetus* pseudocysts were also reported to undergo mitosis [14], adhere cytotoxically to vaginal epithelial cells [10, 19], and form multinucleate bodies from which new trophozoites can bud off [20]. In parallel*, T. vaginalis* pseudocysts were also seen to occur at low frequency (∼5% of cells) in normal axenic culture [14, 21] but at higher frequencies after temperature change and iron depletion [14, 22], and to undergo mitosis [14]. They were also observed in clinical isolates [11, 23]. Hussein and Atwa [24] reported that intravaginal inoculation of mice with *T. vaginalis* pseudocysts produced infection Crucially, two independent studies reported reversion of induced *T. vaginalis* pseudocysts to trophozoites after returning them to normal media [11, 22].

Pseudocysts have also been reported in studies of trichomonad programmed cell death. Benchimol [25] distinguished between apoptosis, programmed autophagy, programmed necrosis, and classical necrosis in protozoan parasites, the last being a ‘fatal accident’ rather than ‘suicide’. Markers of programmed cell death (e.g. apoptosis) in eukaryotes include cell shrinkage, membrane blebbing, altered plasma membrane permeability, exposure of phosphatidylserine (PS) on the cell surface, loss of mitochondrial integrity, chromatin condensation and characteristic DNA fragmentation and protein cleavage [26], eventually leading to necrotic cell membrane rupture. Programmed cell death of unicellular parasites has been extensively studied in *Leishmania,* trypanosomes, and *Plasmodium*, which have conventional mitochondria, but it is less well understood in those which lack true mitochondrial such as trichomonads and *Giardia* [27]. Chose *et al.*, [28] found abundant ‘apoptosis-like bodies’ in exhausted *T. vaginalis* cell cultures (typically ∼72 hr growth in Diamond’s media) and in cultures treated with apoptosis-inducing drugs; these are described morphologically as round with condensed nuclei, and lacking flagellar motion. They have intact plasma membranes and present surface PS after treatment with the apoptosis inducer staurosporine. An ‘irreversible’ form of *Tr. foetus* pseudocyst was induced by hydrogen peroxide treatment and presumed to presage cell death [29].

Silva de Oliveira *et al.* observed both pseudocysts and apoptotic cells in *T. vaginalis* cultures treated with certain metal-metronidazole derivatives [30]. Questions of what advantage suicide has for unicellular organisms, e.g., how ‘altruism’ (as understood to favor close genetic kin) could apply, hang over all such findings [31].

*T. vaginalis* is typically axenically cultured *in vitro* in Diamond’s Trypticase-Yeast extract-Maltose (TYM) media supplemented with horse serum and iron, with pH adjusted to 6.2, based on a 1957 protocol [32]. While TYM culture fosters trophozoite proliferation, it does not emulate the physiological conditions and microbial dynamism of the human vagina. Studying *T. vaginalis* in an *in vitro* context dissimilar from its *in vivo* environment could produce data that do not reflect real-world stresses encountered by the parasite. A major difference between *T. vaginalis* growth *in vivo* and typical *in vitro* conditions is pH. A healthy reproductive-age vagina ranges in pH from 2.8 to 4.2 (average 3.5) [33], while TYM is typically adjusted to pH 6–7, a range the human vagina rarely reaches, if ever. During menses, vaginal pH can increase to around 5 (coinciding with worsening trichomoniasis symptoms [4]), but this is still more acidic than TYM media. Vaginal pH can also increase to around 5 during perimenopause and menopause [34]. Interestingly, Beri *et al*., [11] showed that *T. vaginalis* trophozoites convert to pseudocysts within 24 hours of culture in media at physiological vaginal pH (pH 3–5).Similar to changes in vaginal pH *in vivo*, vaginal iron availability also changes during perimenopause [35]. Iron is essential for *T. vaginalis* metabolism and a recent longitudinal study by Kim *et al*. [35] showed that premenopausal low levels of vaginal ferritin and serum iron sharply increase during perimenopause. Iron depletion has been used by multiple labs to achieve nearly complete conversion of trophozoites to pseudocysts [22, 36].

While there have been several *T. vaginalis* trophozoite transcriptome studies (for example [37, 38]), there have been a few on *T. vaginalis* pseudocysts. These include Cheng *et al*., [39] who showed that expression of oxidoreductases and glutamate dehydrogenase increased in iron depletion-induced pseudocysts, and that they accumulated nitric oxide, and an analysis by Huang *et al*., [40] of the transcriptome of *T. vaginalis* cultured under glucose restriction that reported that energy metabolism-related genes were downregulated and autophagy was induced. A recent metabolomics study also provided evidence for cellulose polymerization and glycogen degradation in *T. vaginalis* pseudocysts induced in iron-depleted conditions compared to trophozoites [36]. However, these previous studies focused on comparing trophozoite datasets against a single pseudocyst time point dataset, without investigating the profile of the pseudocyst over time.

Here, we present a more comprehensive investigation of the changes exhibited by *T. vaginalis* parasites as they transition from trophozoites to pseudocysts. We first compared the efficacy of acidic media and iron depletion methods for inducing pseudocysts, finding that media adjusted to the physiological pH of a pre-menopausal vagina kills *T. vaginalis* and is not an appropriate method of inducing live pseudocysts. We then present our findings based on flow cytometry and transcriptomic analyses that *T. vaginalis* pseudocysts are a viable, metabolically quiescent cell stage distinct from dead parasites. We find that pseudocysts maintain plasma membrane integrity across days and expose phosphatidylserine on their surfaces. We also demonstrate that while returning iron depletion-induced pseudocysts to iron-replete TYM media is insufficient to trigger reversion to the trophozoite stage, any residual trophozoites from incomplete pseudocyst induction will proliferate under these conditions. These findings emphasize the necessity of isolating pure pseudocyst populations to ensure the accuracy of future reversion studies. Finally, pseudocysts induced by iron depletion maintain intact RNA for at least five days. While the majority of the genes show decreased expression, there are hundreds of protein-coding genes with increased expression that are the same across several days, suggesting a concerted stress response mechanism. Additionally, we find evidence for the transcription of pseudocyst-specific genes that are not expressed in trophozoites.

## Results

### Iron depletion, but not low pH, is a reproducible and physiologically relevant method of inducing pseudocysts

During routine treatment of *T. vaginalis in vitro* cultures of clinical isolates with the antifungal drug Nystatin to clear *Candida* spp. symbionts, we observed that many *T. vaginalis* parasites had formed pseudocysts (**Fig 1A**). In order to develop a more standardized, reproducible, and physiologically relevant method of pseudocyst generation in the lab, we reviewed published methods for experimentally inducing pseudocysts and focused on two with physiological relevance. The method described in Beri *et al*., cultures *T. vaginalis* trophozoites in acidic media for 48 hours cultures [11]; while Dias-Lopes *et al.,* use iron depletion to induce pseudocyst formation in *T. vaginalis* cultures [22].

**Figure 1.**
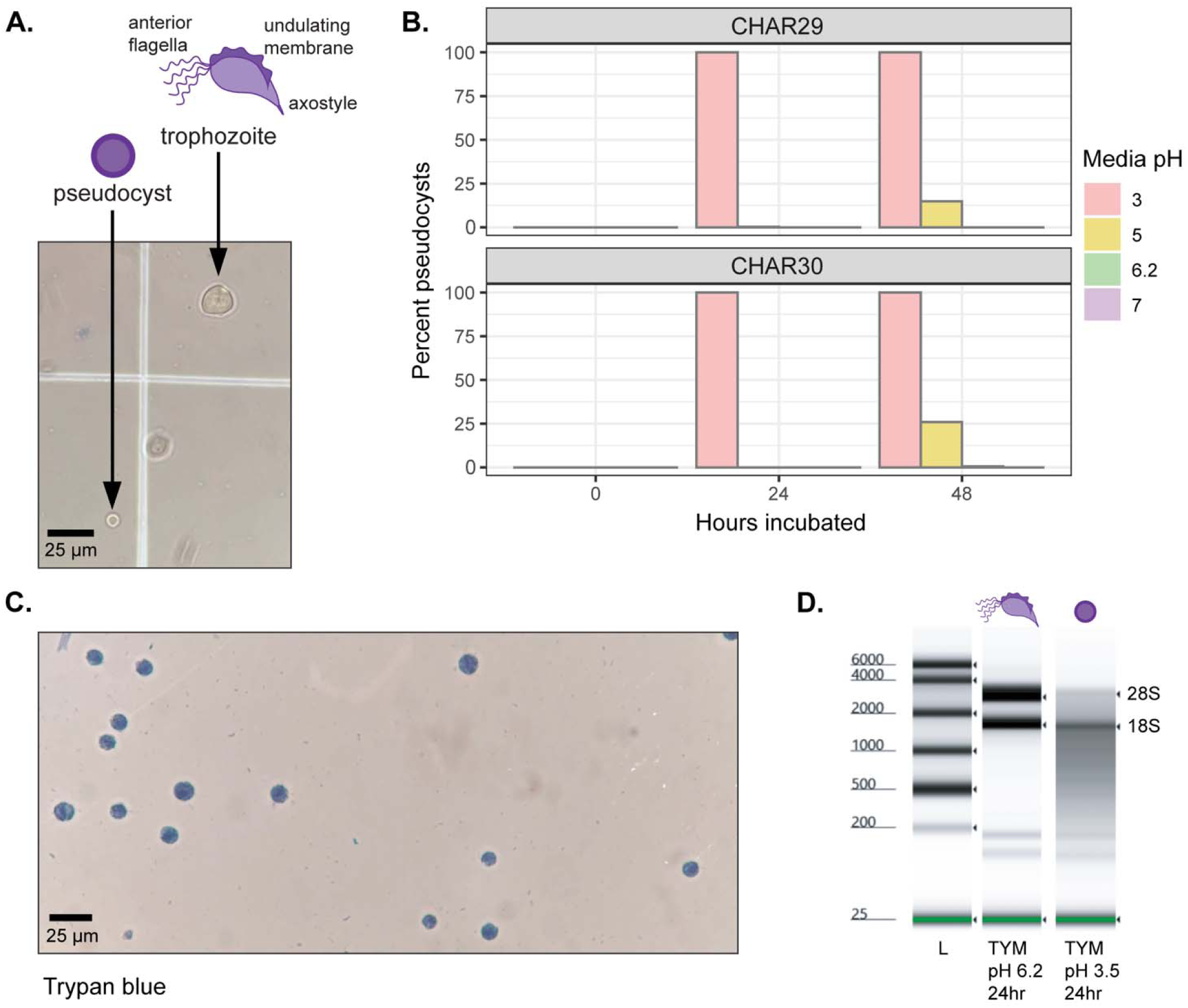
Acidic media induction of *T. vaginalis* pseudocysts and parasite death. (A) Bottom: light microscope image of a *T. vaginalis* trophozoite, and a pseudocyst induced by Nystatin treatment. Top: cartoon representation of typical pseudocyst and trophozoite morphologies. (B) Bar chart showing conversion to trophozoites to pseudocysts measured every 24 hours for *T. vaginalis* strains CHAR29 (top) and CHAR30 (bottom) incubated in media adjusted to pH 3.0, 5.0, 6.2 and 7.0. Media at pH 3.0 (pink) induced complete conversion to pseudocysts for both strains. (C) Representative light microscope field of strain CHAR29 parasites after 24 hours in pH 3.5 media stained with trypan blue. (D) TapeStation electropherogram of *T. vaginalis* CHAR29 RNA extracted from parasites incubated in pH 6.2 TYM (trophozoite cartoon) and pH 3.5 TYM (pseudocyst cartoon) for 24 hrs.

We first tested the low pH method growing two *T. vaginalis* strains in TYM media adjusted to pHs 3.0, 5.0, 6.2, and 7.0 media for 24 and 48 hours. As shown in **Fig 1A**, after 24 in pH 3.0 media all cells from both strains formed pseudocyst-like cells that were rounder and smaller than trophozoites. In pH 5.0 media, 14.9% of strain CHAR29 and 25.9% of strain CHAR30 formed pseudocyst-like cells by 48 hours while we did not detect these cells in pH 6.2 or pH 7.0 (**Fig 1B**). While these morphological changes were consistent with pseudocysts, all of them were stained by trypan blue dye indicating that these are non-viable cells (**Fig 1C**). Consistent with the trypan blue staining, RNA extracted from these parasites was fully degraded suggesting that the round cells we obtained from acidic cultures were dead parasites rather than true pseudocysts (**Fig 1D**).

Next, we investigated iron depletion as a method of pseudocyst induction. We cultured the same two *T. vaginalis* strains in iron depleted media (TYM-DIP) and we then collected morphometric and growth data following the methods described in Dias-Lopes *et al.* [22]. As expected for parasites grown in complete TYM media, cells grew exponentially for about 36h until they hit carrying capacity and the population collapsed. In contrast, parasites grown in TYM-DIP media remained at low numbers for the entire experiment (**Fig 2A, B**). These population dynamics closely reproduce the data from the original report [22]. Microscopic analyses of cells cultured in TYM-DIP for 24h, revealed that most parasites adopted the round and smaller morphology typical of pseudocysts and by 36 hours all cells showed this phenotype (**Fig 2C, D**). In contrast to cells grown in acidic media, after iron depletion cells remained viable as they did not stain with trypan blue (**Fig 2E**) and their RNA remained stable for at least 96 hours of culture in TYM-DIP media (**Fig 2F**).

**Figure 2.**
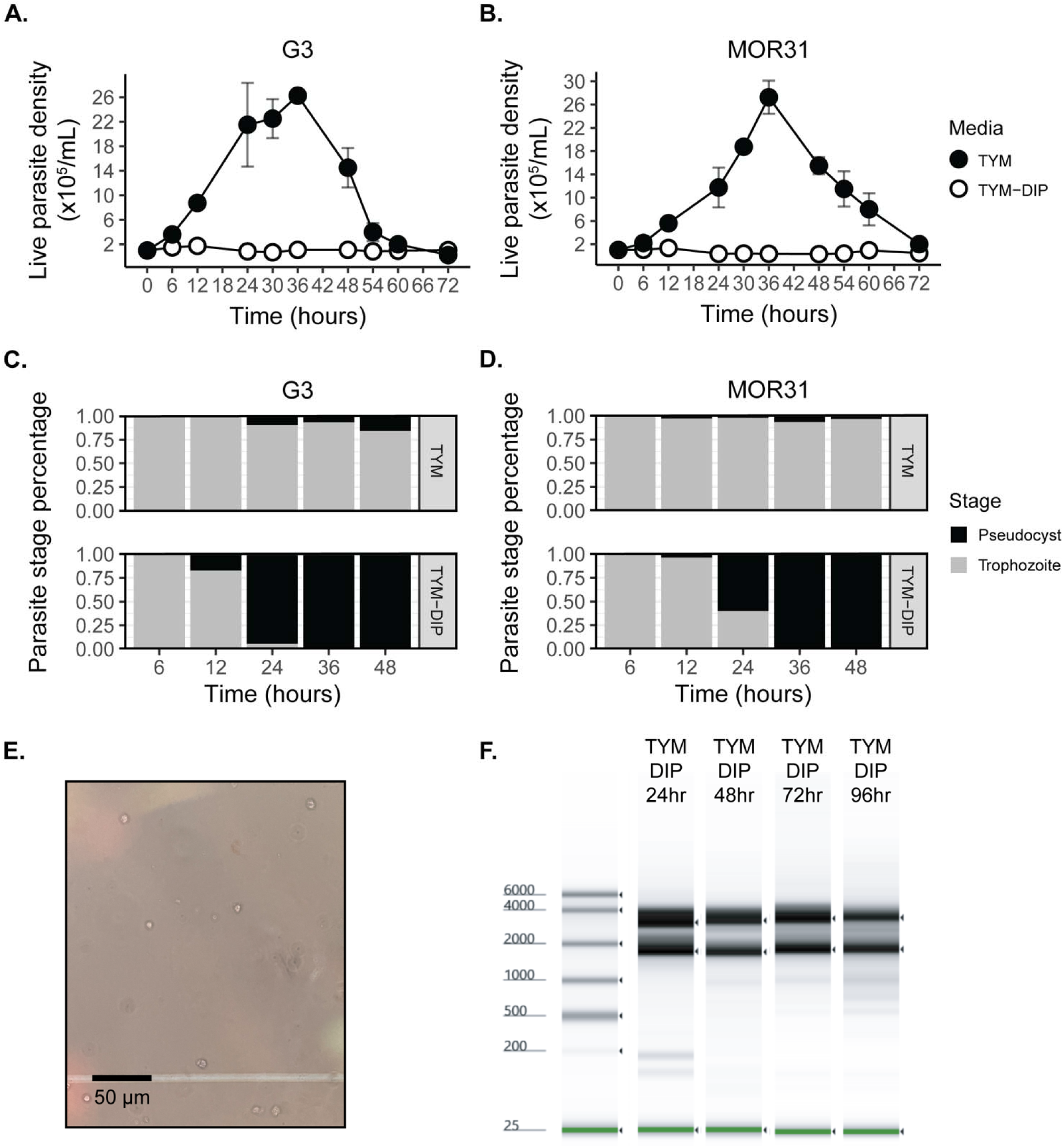
Iron depletion induction of *T. vaginalis* pseudocysts and cell viability. (A, B). Growth curves of *T. vaginalis* strains G3 and MOR31 cultured in TYM media (black circles) or TYM-DIP media (white circles) for 72 hours at 37°C. Experiments were performed in triplicate and results are shown as medians of live parasite density ± SD. (C, D) Proportions of *T. vaginalis* life stages present in cultures of two strains over 48 hours of incubation in TYM (top) or TYM-DIP (bottom) media. (E) Light microscopy field of iron depletion-induced pseudocysts stained with Trypan blue (40x objective). (F) TapeStation electropherogram of RNA extracted from strain G3 parasites incubated in TYM-DIP for 24 hrs, 48 hrs, 72 hrs, 96 hrs showing high integrity and little degradation.

Combined, these data showed that iron depletion induced viable pseudocysts with stable RNA in a reproducible manner. We thus used this method for pseudocyst generation in all subsequent experiments.

### Flow cytometry experiments confirm distinct parasite cell states and pseudocyst cell membrane integrity

Next, we used flow cytometry to further profile the phenotype of *T. vaginalis* pseudocysts. Initial experiments of two *T. vaginalis* strains cultured under iron depletion conditions for 0, 1, 2, 3, and 7 days identified two distinct populations: predominantly trophozoites on Day 0 and predominantly pseudocysts on Days 1, 2, 3 and 7 (**S1 Fig**). To assess the metabolic activity of pseudocysts, parasites that were incubated in TYM-DIP were stained with fluorescein diacetate (FDA) and analyzed by flow cytometry. FDA fluorescence indicates esterase activity and is used as an indicator of general cell metabolic activity. Day 0 trophozoites had high FDA fluorescence, but the signal rapidly disappeared indicating that pseudocysts were not metabolically active (**Fig 3A**).

**Figure 3.**
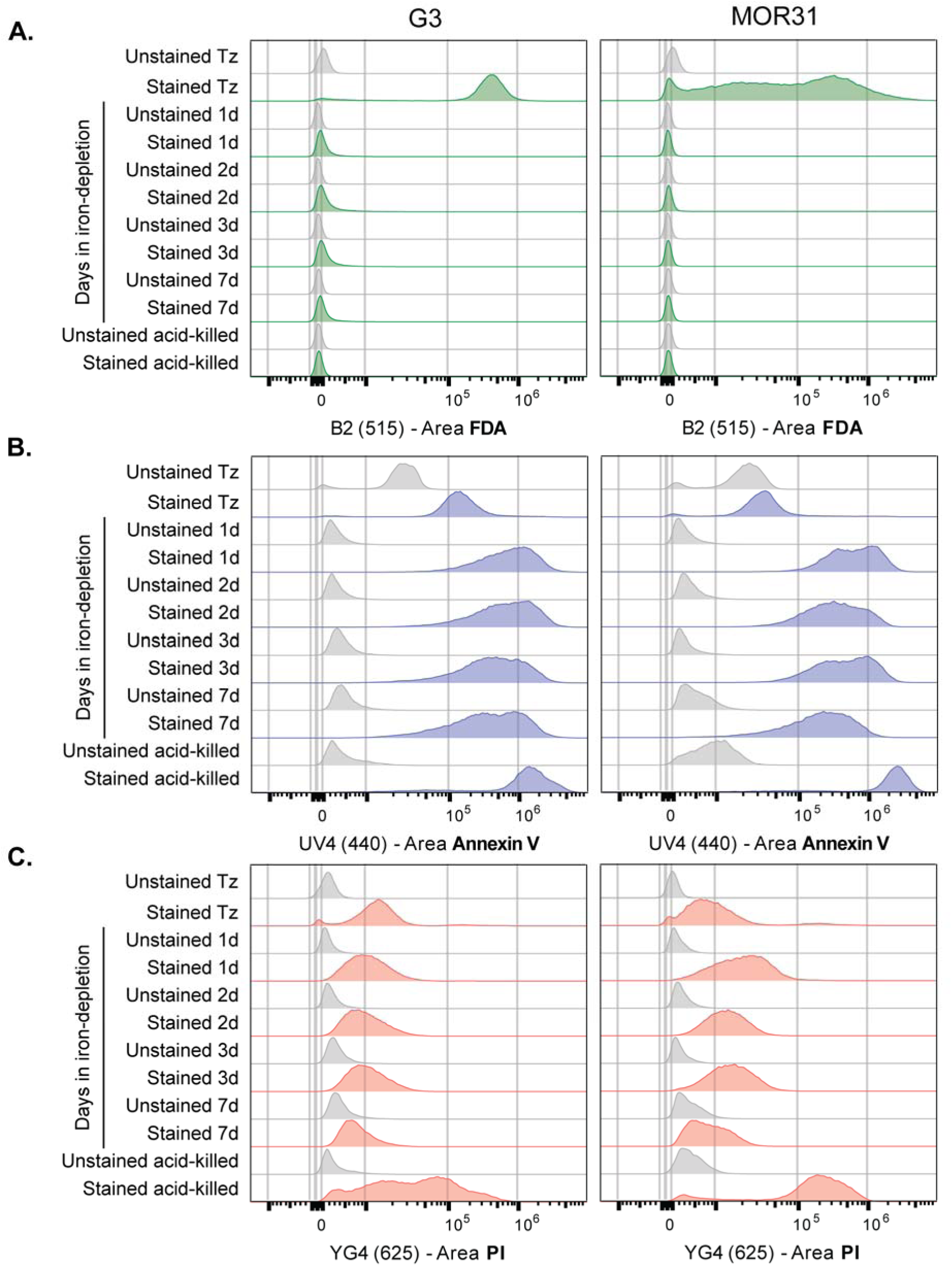
Flow cytometry of *T. vaginalis* trophozoites and pseudocysts. Histograms of (A) fluoresceine diacetate (FDA), (B) Annexin V, and (C) Propidium Iodide (PI) fluorescence of two *T. vaginalis* strains G3 and MOR31 incubated in iron depleted media for 0d (Tz), 1d, 2d, 3d, and 7d. Stained data are colored and unstained control data are grey. Numbers next to the stain names refer to the specific fluorescence detection channel used. Tz: trophozoite.

Since these pseudocysts appear metabolically inactive but viable, we asked if they showed the loss of additional energy-dependent processes. In healthy eukaryotic cells, energy-dependent flippases and floppases maintain distinct asymmetric distributions of phospholipids in the two leaflets of the plasma membrane. Under different conditions of stress, cells activate ATP-independent lipid scramblases that passively equalize the lipid distribution exposing for example phosphatidylserines on the outer surface, whereas these lipids are normally only exposed to the cytosolic side of the membrane. To assess whether pseudocysts expose phosphatidylserines, we stained them with Annexin V that specifically binds to these phospholipids. At the same time, we used propidium iodide (PI) to assess cell viability. Our experiments revealed that pseudocysts had a strong Annexin V signal (**Fig 3B**), and yet they remained viable for at least 7 days (**Fig 3C**). These results are consistent with the loss of metabolic activity (as determined via FDA) while retaining membrane integrity as these cells are not stained by PI nor by trypan blue. In contrast, trophozoites incubated in acidic media at pH 3.5 for 24 hours, completely lost their membrane integrity.

### Iron depletion-induced pseudocysts do not revert to trophozoites upon return to iron-replete conditions

We assessed reversion of iron depletion-induced pseudocysts upon return to standard TYM media in two *T. vaginalis* strains. Parasites were incubated in TYM-DIP for 1 day (producing a partially converted population consisting of trophozoites and pseudocysts) or 2 days (producing a fully converted population of only pseudocysts), then transferred to standard TYM media and observed after 1 day or 3 days. In both strains, the pure pseudocysts cultures incubated in TYM-DIP for 2 days did not show evidence of trophozoites re-emerging after 1 day or 3 days of return to TYM (**S2 Fig**, **S3 Fig**). The partially converted cultures incubated in TYM-DIP for 1 day showed increased trophozoite populations after 24 hours of return to standard TYM media. This trend continued after 3 days, indicating that residual trophozoites that survived iron depletion rapidly proliferated upon reintroduction into iron-replete TYM.

To further investigate whether iron depletion-induced pseudocysts reverted to trophozoites upon return to standard TYM media, we separated residual trophozoites from pseudocyst populations using FDA fluorescence and fluorescence-activated cell sorting (FACS), with pseudocysts classified as FDA-positive and trophozoites as FDA-negative. Log growth trophozoites from two *T. vaginalis* strains were incubated in TYM-DIP for 0, 1, 2 days, and four populations were sorted per strain: Day 0 pseudocysts, Day 0 trophozoites, Day 1 pseudocysts, and Day 2 pseudocysts (**Fig 4A**, **C**). After cell sorting, the four populations were incubated in TYM for three days to allow for any reversion or residual trophozoite proliferation to occur and then visualized with flow cytometry again (**Fig 4B**, **D**).

**Figure 4.**
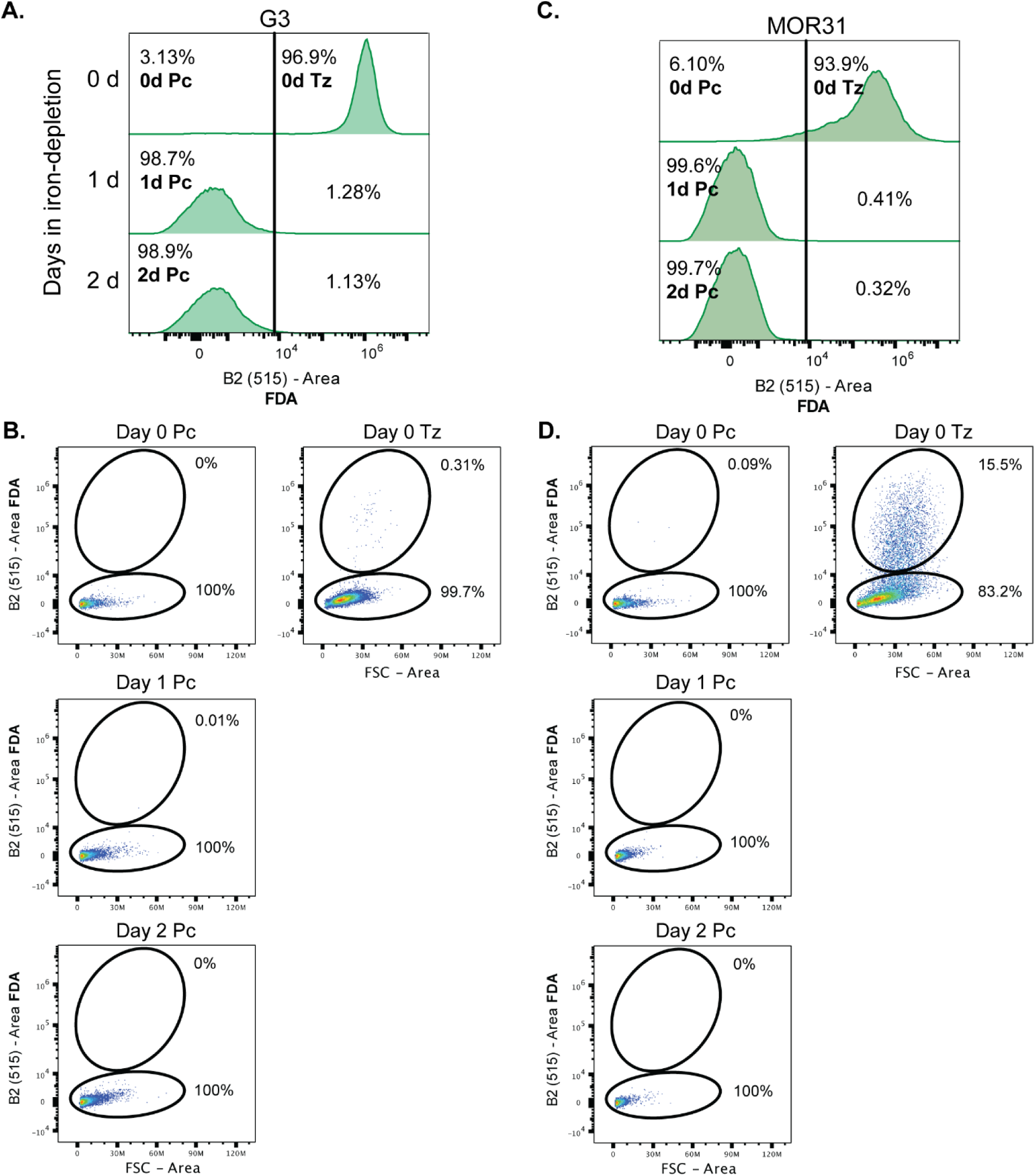
Iron depletion-induced *T. vaginalis* pseudocysts do not revert to trophozoites when returned to iron-replete media. Histograms of the FDA fluorescence of *T. vaginalis* strains (A) G3 and (C) MOR31 incubated in TYM-DIP for 0-2 days. The vertical line is the fluorescence sorting threshold and the populations that were sorted are indicated in bold. Numbers next to the fluorescent dye FDA name refer to the specific fluorescence detection channel used. Flow cytometry plots of the sorted populations of strains (B) G3 and (D) MOR31 incubated in iron-replete TYM media for three days. FDA fluorescence (y-axis) is shown plotted over area (x-axis). Ovals delineate regions of interest, and percentages of the populations in the trophozoite (top ovals) or pseudocyst (bottom ovals) are shown on each plot. Abbreviations: Pc, pseudocyst; Tz, trophozoite.

The trophozoites sorted from Day 0 populations showed proliferation and subsequent decline (due to population decline after nutrient depletion) after three days in TYM, although some viable cells nonetheless remained. None of the other populations had more than three events in the trophozoite gate, and most had none. Thus, there was no evidence of population-level reversion of pseudocysts to trophozoites upon return to iron-replete standard TYM media.

Finally, we used “spent” media from actively growing trophozoites to test whether any potential trophozoite-secreted signaling factors might trigger reversion of pseudocysts back into trophozoites. Iron depletion-induced *T. vaginalis* pseudocysts were resuspended in four kinds of media: (1) standard TYM, (2) TYM-DIP, (3) filtered media taken from log-phase growing parasites in TYM (“spent media”), and (4) a 50/50 mix of spent media and fresh TYM (called “half spent media”). The pseudocysts were incubated at 37°C for 41 days, counted, and viability determined using trypan blue (**S4 Fig**). Parasite viability declined in all media types over time, and trophozoites were not seen to re-emerge. Notably, even after 41 days, 9% of the parasites in TYM-DIP and 12% of the parasites in TYM remained alive with intact membranes.

### Pseudocysts are transcriptionally distinct from trophozoites and appear to be metabolically quiescent

With evidence to support that pseudocysts are not dead cells, we next explored their transcriptional profile. We generated RNA-seq libraries from two *T. vaginalis* strains incubated in TYM-DIP for 0 to 5 days, with each day represented as three biological replicates, and sequenced them on an Illumina NovaSeq 6000 generating paired-end reads. An average of 27,987,377 reads were generated per library after adapter trimming. A principal component analysis (PCA) of the RNA-seq datasets showed a strong effect along the PC1 axis, with Day 0 datasets (trophozoites) clustered at one end and Day 2–5 datasets (pseudocysts) clustered together at the opposite end (**Fig 5**). Day 1 samples (samples with a mix of trophozoites and pseudocysts), clustered in between. Triplicate RNA-seq libraries for each day were tightly clustered, indicating a high degree of experimental reproducibility and reliability within biological replicates. In both *T. vaginalis* strains there were hundreds to thousands of genes with differential expression in pseudocysts compared to trophozoites (**S1 Table**).

**Fig 5.**
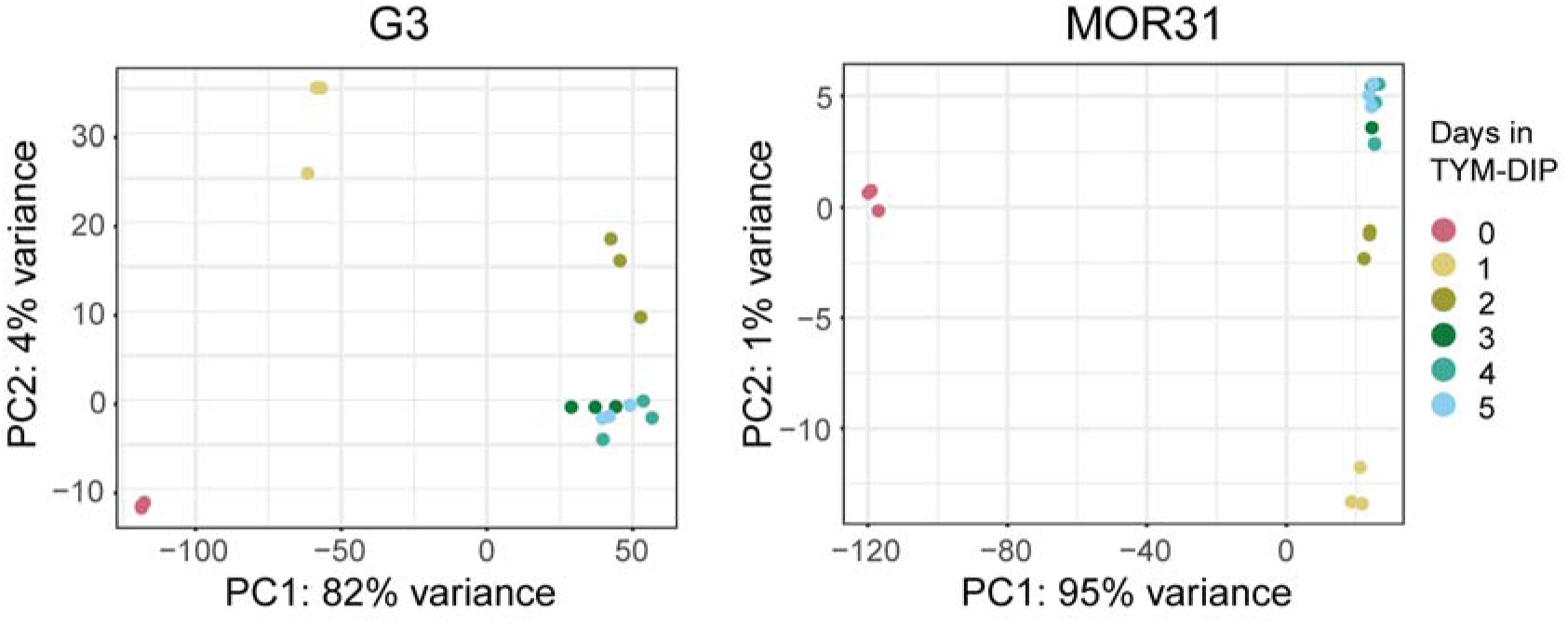
PCA plots of gene expression data of two *T. vaginalis* strains during iron depletion. PCA plots of RNA-seq data from two *T. vaginalis* strains (left: strain G3; right: strain MOR31) incubated in TYM-DIP media for 0-5 days; Day 0 (trophozoites), Day 1 (mixed trophozoites and pseudocysts), and Days 2–5 (pseudocysts). Three data points are shown for each condition representing the biological triplicate libraries generated for each day. PC1 is plotted on the x-axis representing the greatest amount of variance in both strains, and PC2 is plotted on the y-axis representing the second greatest amount of variance in both strains.

Differential gene expression analysis was also performed on the RNA-seq data from each of the two *T. vaginalis* strains G3 (**Fig 6**) and MOR31 (**S5 Fig**) and showed similar results for both strains. We generated a pseudocount-adjusted gene count matrix (see **Materials and methods**) in order to construct volcano plots to illustrate the differentially expressed genes in pseudocysts on each day of iron depletion compared to trophozoites (**Fig 6A**, **S5A Fig**). The number of genes with increased expression remained consistent over time in both strains of *T. vaginalis*, while the number of genes with decreased expression increased over time in both strains, most obviously in strain MOR31 (**Fig 6A**, **S5 Fig**). Intersecting the genes at Days 2, 3, and 4 in each expression category (increased or reduced) showed that, within each strain, the same genes showed increased expression from one day to the next (**Fig 6B**, **S5B Fig**), i.e., gene expression changes were not random. There was less, but still significant, overlap in the genes with reduced expression over time (**Fig 6C**, **S5C Fig**). Of the 364 protein-coding genes in strain G3 with increased expression in common amongst the three days, 61.5% of them had an ortholog among the 3-day-overlapping MOR31 protein-coding genes with increased expression. Of the 529 G3 protein-coding genes with decreased expression in common amongst the three days, 28.4% of them had an ortholog among the 3-day-overlapping MOR31 protein-coding genes with decreased expression. A SANT/Myb domain-containing gene (TVAGG3_0200340 and TVAGMOR31_0200340) had the greatest differential gene expression in both *T. vaginalis* strains. This gene had no expression in trophozoites for either strain but was expressed in the pseudocyst samples for both (**Fig 6D**, **S5D Fig**). Finally, we investigated the expression of several key experimentally validated *T. vaginalis* autophagy genes required for autophagosome assembly [41–43]: Atg3 (TVAGG3_1001600/ TVAGMOR31_1001600), Atg8a (TVAGG3_0691340/TVAGMOR31_0691340), and Atg8b (TVAGG3_0430750/TVAGMOR31_0430750), and found that they had higher expression in pseudocysts compared to trophozoites (**Fig 6E**, **S5E Fig**).

**Fig 6.**
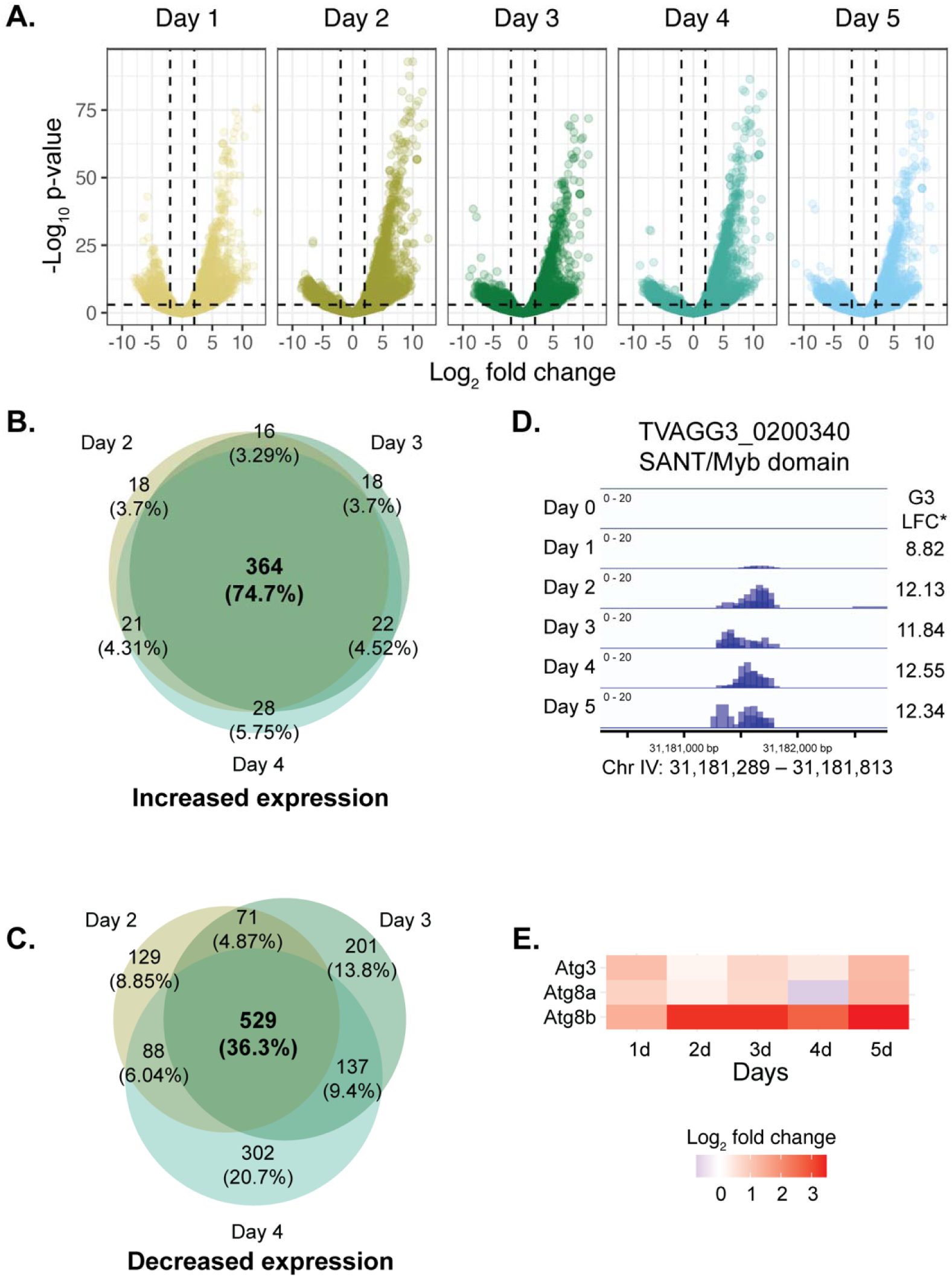
Differential expression analysis of pseudocyst genes with increased expression compared to trophozoites in *T. vaginalis* strain G3. (A) Volcano plots showing differential gene expression of protein-coding genes in *T. vaginalis* G3 grown in iron-depleted media for 1-5 days compared to day 0 (trophozoites). The x-axes represent the log_2_ fold change and the y-axes represent the log_10_ p-value. Dashed vertical lines are the threshold for genes with decreased expression (log_2_ fold change < -2) and increased expression (log_2_ fold change > 2). Dashed horizontal lines represent a significance threshold of p-value < 0.001. Venn diagrams showing the (B) overlap of protein-coding genes with increased expression and (C) reduced expression on Days 2, 3, and 4 of iron depletion compared to trophozoites. Numbers in bold represent the number of genes and their percent overlap over all three days. (Day 1 and Day 5 data are not shown since Day 1 cultures contained a mixture of trophozoites and pseudocysts, and Day 5 RNA samples had a RIN value less than 9). (D) IGV read coverage of the protein-coding gene with the highest expression in pseudocysts compared to trophozoites, TVAGG3_0200340. (E) Heat map of log_2_ fold change values of autophagy genes Atg3, Atg8a, and Atg8b in *T. vaginalis* strain G3 in culture for 1-5 days in iron depletion compared to trophozoites.

### Deconvolution analysis identifies different *T. vaginalis* cell types induced by iron depletion

We next profiled the transcriptomes over time to understand the changes occurring to *T. vaginalis* parasites during iron depletion. We used the cell type deconvolution program CDSeq, which models bulk RNA-seq read counts using a probabilistic mixture model, to identify and classify the different *T. vaginalis* cell types revealed across time [44]. A gene expression profile for *T. vaginalis* G3 trophozoites was first generated to define the control cell type (trophozoites) as a parameter for the model. We compared three independent RNA sequencing experiments of trophozoites performed in triplicate and identified 26,356 genes with a transcripts-per-million >0 across all experiments and replicates. Further filtering the *T. vaginalis* RNA-seq gene count matrix to include only genes with an effective length > 1 (the number of possible start sites to which a library fragment can be mapped to within a gene) and a read count > 10 across replicates and timepoints identified 25,535 genes that satisfied these thresholds.

After 1,500 iterations the model converged on eight cell types (**Fig 7A**); no convergence occurred after Day 3 and so subsequent analyses focused on Day 0 through Day 3. The four cell types that did not reach a proportion greater than 0.2 over the four days were considered artifactual (Cell Types 1, 2, 7, 8). Cell Types 4 and 6 are likely transient cell types formed in response to an acute stress on Day 1 and that subsequently disappeared. In the model, one cell type (Cell Type 5) was predominant at Day 0 (60%) but disappeared from the population after one day of iron depletion, while another cell type (Cell Type 3) composed most of the population after Day 2 (59.7%) and Day 3 (72.6%) of iron depletion. This suggests that Cell Type 5 were trophozoites and Cell Type 3 were pseudocysts (the percentages of each cell type on each day are provided in **S2 Table**). We performed differential gene expression analysis on Cell Type 3 (pseudocysts) and found that this cell type had significantly reduced gene expression on Days 1, 2, and 3 compared to Day 0 (**Fig 7B**).

**Fig 7.**
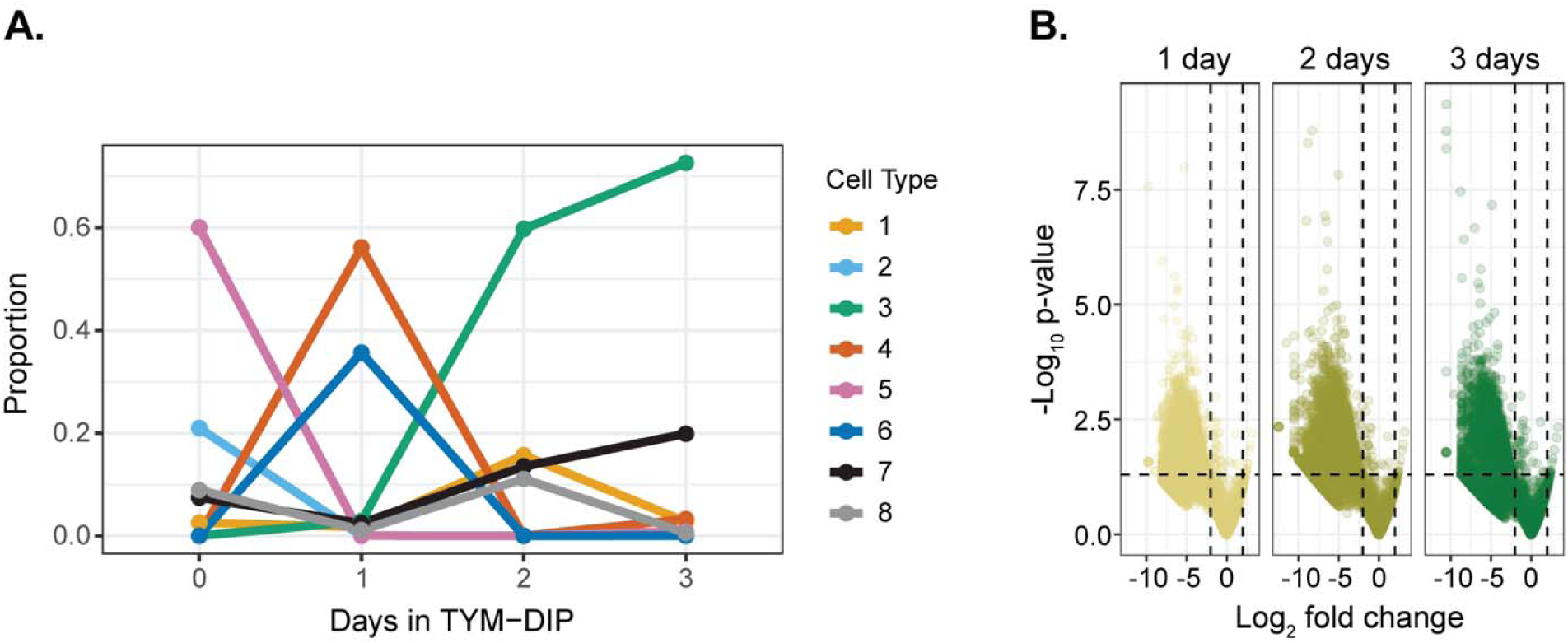
*T. vaginalis* cell types identified under iron depletion conditions. (A) Proportion of different cell types (y-axis) differentiated over days incubated in iron depleted media (x-axis). (B) Volcano plot showing gene expression of *T. vaginalis* Cell type 3 (pseudocysts) for 1-3 days in iron depletion, graphed as p-value (y-axis) over log_2_ fold change (x-axis). Horizontal dashed line represents a p-value of 0.05 and vertical dashed lines represent log_2_ fold change values of -2 and 2. The results shown are limited to Days 0–3 since the model failed to converge when Days 4 and 5 were included.

## Discussion

The sexually transmitted parasite *T. vaginalis* has two well-known cell forms: a free-swimming, flagellated trophozoite and an amoeboid form adhered to host epithelia. A third morphology, the pseudocyst, has been described, but it is unclear whether this is a quiescent stage capable of facilitating persistent infections, or a degenerate form indicating cell death. Here we describe flow cytometry, fluorescent activated cell sorting, whole transcriptome, and cell type deconvolution experiments that reveal pseudocysts can be a viable cellular state in *T. vaginalis* distinct from cell death, but can also present characteristics of programmed cell death.

Iron depletion has been used by several labs to achieve near-complete conversion of trophozoites to pseudocysts [22, 36]. We corroborated iron-depleted *T. vaginalis* growth curves and morphology results presented by Dias-Lopes *et al.,* [22] in two strains *T. vaginalis* G3 and MOR31. Unlike with the acid-induction method introduced by Beri *et al.,* [11], iron depletion-induced pseudocysts maintained their membrane integrity, as indicated by their impermeability to trypan blue and propidium iodide. Their RNA remained intact across four days of iron depletion and did not undergo the degradation we observed in acidic media-induced pseudocysts. Along with shifts in vaginal pH, iron availability also changes with the onset of menopause, increasing during perimenopause [35].

We detected phosphatidylserine (PS) on pseudocyst surfaces and observed the cell membrane to remain intact for at least seven days. Surface PS and membrane integrity together are two characteristics of apoptosis-like programmed cell death [45]. To phagocytes, external PS is a marker of an apoptotic cell, not of a ‘threat’. Engulfing such cells is thought to induce signaling that inhibits an inflammatory response [26]. For *T. vaginalis*, external PS-positive pseudocysts thus might serve as immunological decoys, allowing trophozoites to maintain an infection while avoiding a broader inflammatory response. An investigation into the response of host immune cells to pseudocysts compared to trophozoites is a ripe area for future research. Transcriptional data from host cells would reveal whether pseudocysts trigger the same inflammatory pathways as trophozoites or if they participate in a form of immune evasion. And transcriptional studies of co-cultures of immune cells with parasites could provide insight concerning gene expression during *in vivo* infections.

But how would *T. vaginalis* pseudocysts “know” how to sacrifice themselves? Programmed cell death constitutes suicide by a unicellular organism, something hard to reconcile with Darwinian survival imperatives [31]. One proposal is that such deaths regulate parasite density when high density could endanger parasite or host survival, although such ‘apoptosis as a social trait’ is more plausible for genetically clonal infections and only if parasites can communicate to each other [46]. In what has become an exciting field of study, *T. vaginalis* parasites can communicate with each other via extracellular vesicles (EVs; [47, 48]): could EVs also be a means by which they regulate their cell death? Applying single-cell RNA sequencing (scRNA-seq) to *T. vaginalis* parasites would allow greater resolution to dissect the heterogeneity of parasite populations, and more specifically, within pseudocyst populations. ScRNA-seq could identify transcriptional markers that distinguish pseudocysts destined for cell death from those that are potentially capable of long-term persistence.

Transcriptomic studies utilizing recently developed vaginal organoid models [49] would enable the integration of physiological stressors like acidic pH and iron restriction into an *in vitro* system that is more representative of clinical *in vivo* conditions. By more accurately reflecting physiological conditions, transcriptomic studies using these models would likely yield data that is more aligned with the reality of *T. vaginalis* gene expression during an infection compared with a traditional axenic culture.

Previous transcription studies focused on comparing trophozoite datasets against a single pseudocyst time point dataset, without investigating whether the pseudocyst transcriptional profile also changes over time [22, 36, 39, 50]. We corroborated the results showing that trophozoites and pseudocysts have distinct transcriptional profiles. However, we found that pseudocyst expression of gene subsets differs over time, temporarily increasing for some genes after one or two days in iron-depleted media likely as an acute stress response. Other genes showed increased expression for several days, while most of the transcriptome had decreased expression. In our two *T. vaginalis* strains, the same gene (TVAGG3_0200340 in strain G3, TVAGMOR31_0200340 in strain MOR31) encoding a protein with a SANT/Myb domain was the most highly expressed in pseudocysts compared to trophozoites. SANT/Myb domains are known to be involved in cell cycle regulation and chromatin remodeling, and a paper by Schwarz *et al.* [51], suggests they are involved in stress response and cell differentiation in plants and apicomplexans. While the role of SANT/Myb domains in *T. vaginalis* has not been studied specifically, they could potentially play a similar role in the transition of trophozoites to pseudocysts. Genes like TVAGG3_0200340/TVAGMOR31_0200340 could serve as diagnostic markers: determining pseudocyst age *in vivo* (i.e., the time since conversion from trophozoites) may provide additional clinical information for patient treatment and the persistent transcription of these genes make them good contenders.

Three *T. vaginalis* autophagy proteins have been experimentally validated and the genes for all three, Atg3 [43], Atg8a [41, 42], and Atg8b [41, 42] showed increased expression in pseudocysts compared to trophozoites in our study. This suggests that autophagy could be a part of the transition into the pseudocyst stage and for survival in unfavorable conditions. Autophagy has been reported as a survival response of *T. vaginalis* cultured in YI-S media, which is referred to as ‘glucose-restricted’ (although it does contain glucose [52]); the study did not assess pseudocyst presence. Prolonged autophagy can result in cell death [31].

The high degree of overlap in genes with increased expression across two, three, and four days of iron depletion, and between strains, further suggests that *T. vaginalis* exhibits a specific and coordinated response to environmental stressors (in this case iron depletion) and that there is a module of genes responsible for the conversion to pseudocysts. In contrast, the smaller overlap of genes with decreased expression across 2–4 days and the growing number of genes with decreased expression over time are consistent with the non-specific degradation of RNA expected as a consequence of extended exposure to a suboptimal environment.

Evidence to date suggests the fate of a trichomonad pseudocyst can be reversion or death, depending on the severity and duration of the stress it formed in response to [25, 29]. However, the evidence is more convincing for *Tr. foetus* than for *T. vaginalis*. Dias-Lopes et al., [22] grew iron-depleted *T. vaginalis* cultures for 48 hours and found that returning them to iron-replete media for 24 hours was sufficient for essentially complete reversion of a pseudocyst population to trophozoites.

However, their induced pseudocyst cultures still contained ∼3% trophozoites [22]. We only observed such population ‘reversion’ in our experiments when insufficiently pure pseudocyst populations, i.e., parasites cultured in TYM-DIP for only one day and consisting of both trophozoites and pseudocysts, were returned to nutrient-rich media. Pure iron depletion-induced pseudocysts we obtained using FACS did not revert, even after days in culture. This suggests that the results of Dias-Lopes et al., might be due to trophozoite contamination. Beri et al., enriched for ‘cyst-like structures’ (CLS) by pelleting 48 hour *T. vaginalis* cultures and resuspending the pellet in cold distilled water, which selectively lyses trophozoites, for several hours [11]. When placed in fresh culture media at physiological temperature, most (>80%) of these CLS externalized flagella by 72 hours, but only a small percentage (∼9%) had fully reverted to live trophozoites by 3.5 days, the limit of observation. In that study, CLS were identified by chitin staining with calcofluor white (CWF), and CLS and trophozoite viability were assessed by their ability to fluoresce when stained with fluorescein diacetate (FDA). In their hands, CWF only stained CLS, but Kneipp et al., [53] reported chitin in apparent trophozoites of *Tr. foetus* and *T. vaginalis* using several chitin markers, including CWF (although the CWF data were not shown); pseudocysts were not mentioned. While Beri *et al.* also observed typical pseudocyst characteristics in their CLS, such as internalized flagella, spherical shape, and immobility, the discrepant CWF results and the small percentage of full reversions suggest that unequivocal reversion of *T. vaginalis* pseudocysts remains to be documented.

Strict cellular quiescence is defined as a reversible non-dividing stage of arrest where cells retain the ability to re-enter the cell-cycle upon receiving a triggering stimulus [54]. The cells exhibit suppressed transcription, minimal translation, and rearranged metabolic pathways that result in low metabolic activity [54]. In the case of *T. vaginalis*, the host vagina presents multiple challenges, particularly prior to menopause, such as high acidity [34], low iron availability [35], and elevated levels of interferon-epsilon secretion [47, 55], all of which may contribute to trophozoite death but permit pseudocyst survival. However, the triggering stimulus for reversion of pseudocysts to trophozoites is yet unknown and our results show that returning iron depletion-induced pseudocysts to iron-replete conditions is not a sufficient trigger. There is a possibility that such a stimulus exists and has not been characterized yet. It may be that *T. vaginalis* pseudocysts require secreted proteins from host cells, from bacterial species of the reproductive tract microbiome, or from a combination of the changing environmental factors of the menopausal and perimenopausal vagina during which trichomoniasis incidence is at its highest. Using mimetic organoid systems [49], or host cell-parasite co-culture methods, may identify what the triggering stimulus for reversion is. Our findings support that *T. vaginalis* pseudocysts are not a degenerate form of the parasite, but a stress-triggered quiescent yet viable cell stage that *T. vaginalis* adopts in adverse conditions. We raise the possibility that pseudocysts play a strategic role in host evasion and infection persistence, although further study of *T. vaginalis* pseudocysts, for example their interaction with host immune cells, is required.

Furthermore, determining if pseudocysts can revert back to trophozoites and identifying the trigger(s) that may cause this will be key to clarifying the role of pseudocysts in *T. vaginalis* pathogenicity.

## Materials and methods

### T. vaginalis strains and in vitro culture

Four *T. vaginalis* strains were used in this work. Initial studies used *T. vaginalis* strains CHAR29 (NYCC29; available through BEI Resources, identifier NR-58892) and CHAR30 (NYCC30; available through BEI Resources, identifier NR-58893) isolated from patients undergoing a pelvic examination at STD clinics in New York City (NYC) in 2008 [56]. All subsequent work used *T. vaginalis* strain G3 (isolated from Kent, United Kingdom, 1973 [57]; available through ATCC identifier ATCC PRA-98) and strain MOR31 (NYCG31; isolated from NYC in 2008 [56]; available through BEI Resources, identifier NR-58898) due to the subsequent availability of high-quality chromosome-grade genome assemblies and annotations for both strains. Isolates were cultured at 37°C in Diamond’s trypticase-yeast extract-maltose (TYM) media supplemented with 10% horse serum, penicillin, and streptomycin, and iron solution composed of ferrous ammonium sulfate and sulfosalicylic acid as described [58].

Acidic TYM media consisted of standard Diamond’s TYM media adjusted to pH 3.5 with hydrochloric acid instead of pH 6.2. Iron depleted media (TYM-DIP) had no iron solution added and was further supplemented with 180 μM of the iron chelator 2.2-dipyridyl (Fisher Scientific). Cultures were assayed for *Mycoplasma* and yeast and treated with mycoplasma removal agent (MRA; BioRad #BUF035) and/or 12.5 µg/mL of Amphotericin B and 10 µL of Nystatin (Sigma-Aldrich, N9150) per 5 mL of culture volume if found to be contaminated.

### Flow cytometry and fluorescence-activated cell sorting

*T. vaginalis* strains G3 and MOR31 cultures grown to log phase were resuspended in TYM-DIP media at 37°C for 1, 2, 3, and 7 days. Log growth phase trophozoites were used as a control for proliferative, live parasites, and log growth phase trophozoites incubated in acidic TYM pH 3.5 for 24 hours were used as a control for dead parasites. On the day of sorting, parasite samples were centrifuged at 4°C for 10 minutes at 3200 RPM, washed once with PBS, and each resuspended in 100 µL of 1x Annexin Binding Buffer (BD Biosciences, #556454). A total of 1 ug/mL of propidium iodide (PI, ThermoFisher Scientific, #P3566), 2.5 ug/mL of fluoresceine diacetate (FDA, Sigma-Aldrich, #F7378-5G), and/or 4 µL of Annexin V conjugated to Alexa-350 (ThermoFisher Scientific, #A23202) were added to each 100 µL sample, incubated in the dark at room temperature for 15 minutes, and then 400 µL of 1X Annexin Binding Buffer added to each sample. Single stained, unstained, and triple-positive stained samples were prepared concurrently. Samples were analyzed on a BD FACSDiscover S8 Cell Sorter with 50,000 P1 events collected for stained samples and 10,000 P1 events collected for unstained controls.

### RNA isolation and RNA-sequencing

Experiments were carried out in triplicate, starting with parasites growing in log phase and subsequently grown in iron depleted media (TYM-DIP) or in acidic TYM media pH 3.5 media for 24 hours. Parasites were centrifuged at 3200 RPM for 10 minutes at 4°C, washed with 1X PBS twice, resuspended in 350 µL of RLT lysis buffer with 1% 2-Mercaptoethanol, and stored at -80°C until RNA isolation. Total RNA was isolated using the Qiagen RNeasy Mini Kit (Qiagen, #74104) including a column DNase treatment using the Qiagen RNase-free DNase kit (Qiagen, #79254). RNA concentration and quality was quantified using a Nanodrop One spectrophotometer and Agilent TapeStation 4200. RNA-seq library preparation used the Illumina TruSeq Stranded mRNA library preparation kit (Illumina, #20020594) with unique barcodes for each sample using IDT for Illumina TruSeq Unique Dual Index primers (Illumina, #20040871). Libraries were pooled and sequenced on an Illumina NovaSeq 6000 with 200 cycles on a S1 flow cell, generating paired-end reads.

### Bioinformatic analysis

Sequencing read quality was assessed using FastQC (v.0.11.9) and MultiQC (v.1.9). Adapter contamination was removed using Trimmomatic (v.0.39). Differential expression (DE) analysis was performed using the Tuxedo protocol [59] and RNA-seq reads aligned to the new chromosome-grade *T. vaginalis* G3 [60] or MOR31 genomes using STAR (v.2.7.6a) [61] with default parameters. BAM files were processed with a custom Python script to remove chimeric alignments as described [62] and further DE analysis was performed using StringTie (v.2.1.6) and the DESeq package [63] (v.1.44.0) in R (v.4.4.0). Many genes had no coverage in the RNA-seq datasets at one or several time points and since these 0 values presented a challenge for visualizing the DESeq results, a pseudocount of 1 was added to each value of the gene count matrix to generate volcano plots for both strains but not included in any other analyses. *T. vaginalis* RNA-seq data were deposited at the Sequence Read Archive (SRA) of the NCBI under BioProject number PRJNA1449513.

CDSeq analysis of *T. vaginalis* G3 and MOR31 RNA-seq datasets used methodologies originally described by Kang *et al.,* [44]. Only genes with 10 or more sequencing reads across replicates were included in the model. A control trophozoite gene expression profile was generated from three independent RNA-seq experiments done in triplicate in October 2022, January 2024, and December 2024.

## Supporting information

**S1 Fig. Flow cytometry dot plots of *T. vaginalis* strains under iron depletion.** (A) Flow cytometry dot plots of *T. vaginalis* strains G3 and (B) MOR31 parasites that were incubated in iron depleted media (TYM-DIP) for 0, 1, 2, 3, and 7 days shown as area (y-axis) over eccentricity (x-axis). Percentages of the populations included in the trophozoite or pseudocyst regions of interest are included on each plot.

**S2 Fig. Iron depleted *T. vaginalis* stages after incubation in TYM.** Flow cytometry dot plots of *T. vaginalis* strain G3 parasites incubated in iron depleted media (TYM-DIP) for 1 (left) or 2 (right) and put back into TYM for 1 day or 3 days shown as area (y-axis) over eccentricity (x-axis). Percentages of the population included in the trophozoite or pseudocyst regions of interest are indicated on each plot.

**S3 Fig. Iron depleted *T. vaginalis* MOR31 stages after incubation in TYM.** Flow cytometry plots of *T. vaginalis* strain MOR31 parasites that were incubated in iron depleted media (TYM-DIP) for 1 day (left) or 2 days (right) and put back into TYM for 1 or 3 days shown as area (y-axis) over eccentricity (x-axis). Percentages of the population included in the trophozoite or pseudocyst regions of interest are indicated on each plot.

**S4 Fig. Resuspension of iron depletion-induced *T. vaginalis* pseudocysts in TYM and spent media.** Percentage of iron depletion-induced live pseudocysts (y-axis) over time in culture at 37°C (x-axis) after being resuspended in different resuspension medias. Parasite viability was assessed using trypan blue staining visualized by light microscopy.

**S5 Fig. Differential expression analysis of pseudocyst genes with increased expression compared to trophozoites in *T. vaginalis* strain MOR31.** (A) Volcano plots showing differential gene expression of protein coding genes in *T. vaginalis* MOR31 grown in iron depleted media for 1-5 days compared to day 0 (trophozoites). The x-axes represent the log_2_ fold change and the y-axes represent the log_10_ p-value. Dashed vertical lines are the threshold for genes with decreased expression (log_2_ fold change < -2) and increased expression (log_2_ fold change > 2). Dashed horizontal lines represent a significance threshold of p-value < 0.001. Venn diagrams showing the (B) overlap of protein coding genes with increased expression and (C) reduced expression on Days 2, 3, and 4 of iron depletion compared to trophozoites. Numbers in bold represent the number of genes and their percent overlap over all three days. (Day 1 and Day 5 data are not shown since Day 1 cultures contained a mixture of trophozoites and pseudocysts, and Day 5 RNA samples had a RIN value > 9). (D) IGV read coverage of the protein-coding gene with the highest expression in pseudocysts compared to trophozoites, TVAGG3_0200340. (E) Heat map of log_2_ fold change values of autophagy genes Atg3, Atg8a, and Atg8b in *T. vaginalis* strain MOR31 in culture for 1-5 days in iron depletion compared to trophozoites.

**S1 Table. Counts of differentially expressed protein-coding genes in *T. vaginalis* strains G3 and MOR31 pseudocysts compared to trophozoites over five days of iron depletion.** Protein coding genes were filtered for genes with a base mean > 100 and a Benjamini-Hochberg adjusted p-value < 0.001. Genes were determined as having increased expression if they had a log_2_ fold change of > 2 and decreased expression if they had a log_2_ fold change < -2.

**S2 Table. CDSeq cell types classified over three days of iron depletion.** CDSeq cell types 1 through 8 identified from *T. vaginalis* gene expression data of Day 0 (trophozoite), Day 1 (trophozoite and pseudocysts), and Days 2 and 3 of incubation in iron-depleted media.

## Acknowledgments

We thank NYU Center for Genomics & Systems Biology GenCore team for advice and access to high-throughput sequencing services, and Dr. Laura Lee for help with flow cytometry experiments. Research reported in this publication was supported in part through the NYU IT High-Performance Computing resources, services, and staff expertise. MS was partially supported by a New York University MacCracken fellowship and National Institutes of Health T32 Training grant award GM132037. We acknowledge the Zegar Family Foundation for their generous support.

## Author Contributions

**Conceptualization**: Jane Carlton, Carlos Carmona-Fontaine

**Data curation:** Francisco Callejas-Hernandez, Jordan Orosco, Steven Sullivan

**Formal analysis**: Mari Shiratori, Francisco Callejas-Hernandez

**Investigation**: Mari Shiratori, Francisco Callejas-Hernandez, Jordan Orosco

**Supervision**: Jane Carlton

**Visualization**: Mari Shiratori

**Writing – original draft**: Jane Carlton, Mari Shiratori, Steven Sullivan

**Writing – review and editing**: Mari Shiratori, Steven Sullivan, Francisco Callejas-Hernandez, Jordan Orosco, Jane Carlton

