## Supplemental Table 1 for "*Trichomonas vaginalis* pseudocysts are a quiescent cell stage characterized by a differentially regulated transcriptome and intact membranes"

S1 Table.

|  | ***T. vaginalis* strain G3** | | ***T. vaginalis* strain MOR31** | |
| --- | --- | --- | --- | --- |
| **Days in TYM-DIP** | **Increased expression** | **Decreased expression** | **Increased expression** | **Decreased expression** |
| 1 | 497 | 493 | 1783 | 1259 |
| 2 | 420 | 817 | 1818 | 1519 |
| 3 | 419 | 938 | 1792 | 1628 |
| 4 | 435 | 1056 | 1820 | 1671 |
| 5 | 424 | 1230 | 1793 | 1700 |
