## Supplemental Table 2 for "*Trichomonas vaginalis* pseudocysts are a quiescent cell stage characterized by a differentially regulated transcriptome and intact membranes"

S2 Table.

|  | **CDSeq Cell type** | | | | | | | |
| --- | --- | --- | --- | --- | --- | --- | --- | --- |
| **Days** | **1** | **2** | **3** | **4** | **5** | **6** | **7** | **8** |
| **0** | 2.575% | 20.995% | 0.005% | 0.006% | 60.039% | 0.010% | 7.456% | 8.915% |
| **1** | 1.716% | 0.032% | 2.859% | 56.138% | 0.038% | 35.655% | 2.440% | 1.122% |
| **2** | 15.692% | 0.000% | 59.728% | 0.001% | 0.000% | 0.002% | 13.500% | 11.076% |
| **3** | 2.734% | 0.002% | 72.620% | 3.259% | 0.859% | 0.048% | 19.884% | 0.595% |
