## Supplementary figures and images for "*Trichomonas vaginalis* pseudocysts are a quiescent cell stage characterized by a differentially regulated transcriptome and intact membranes"

### Supplemental Fig 1

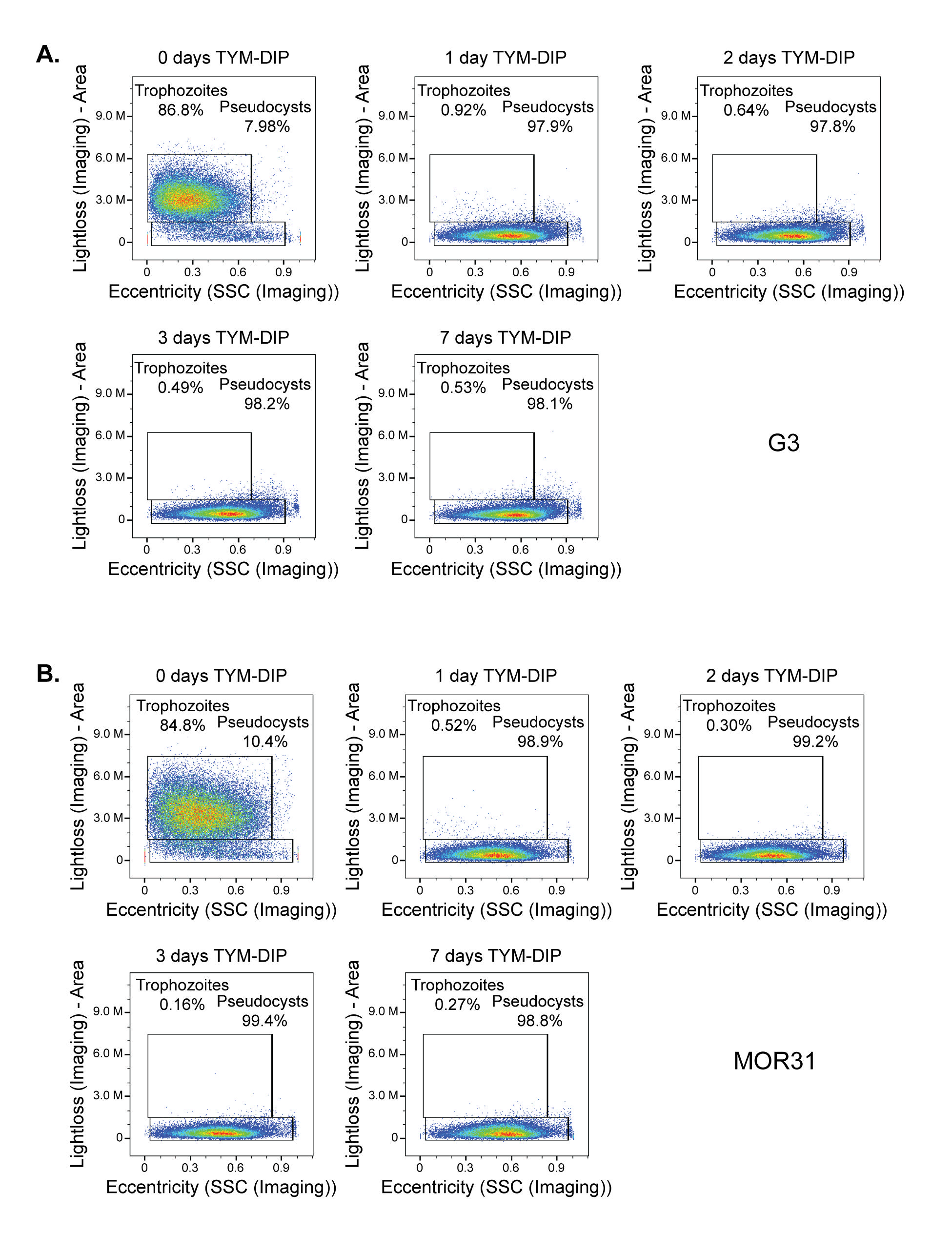

### Supplemental Fig 2

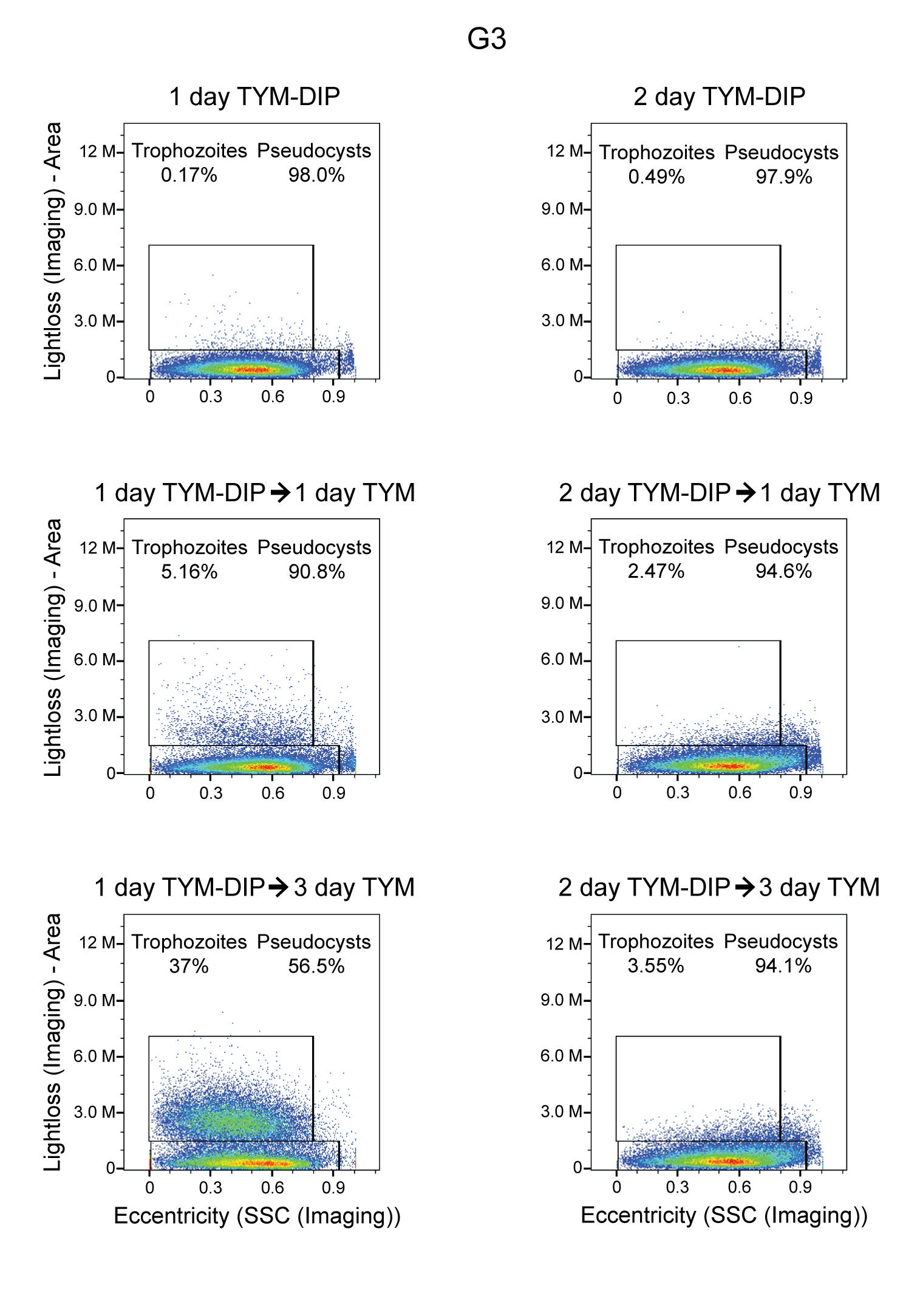

### Supplemental Fig 3

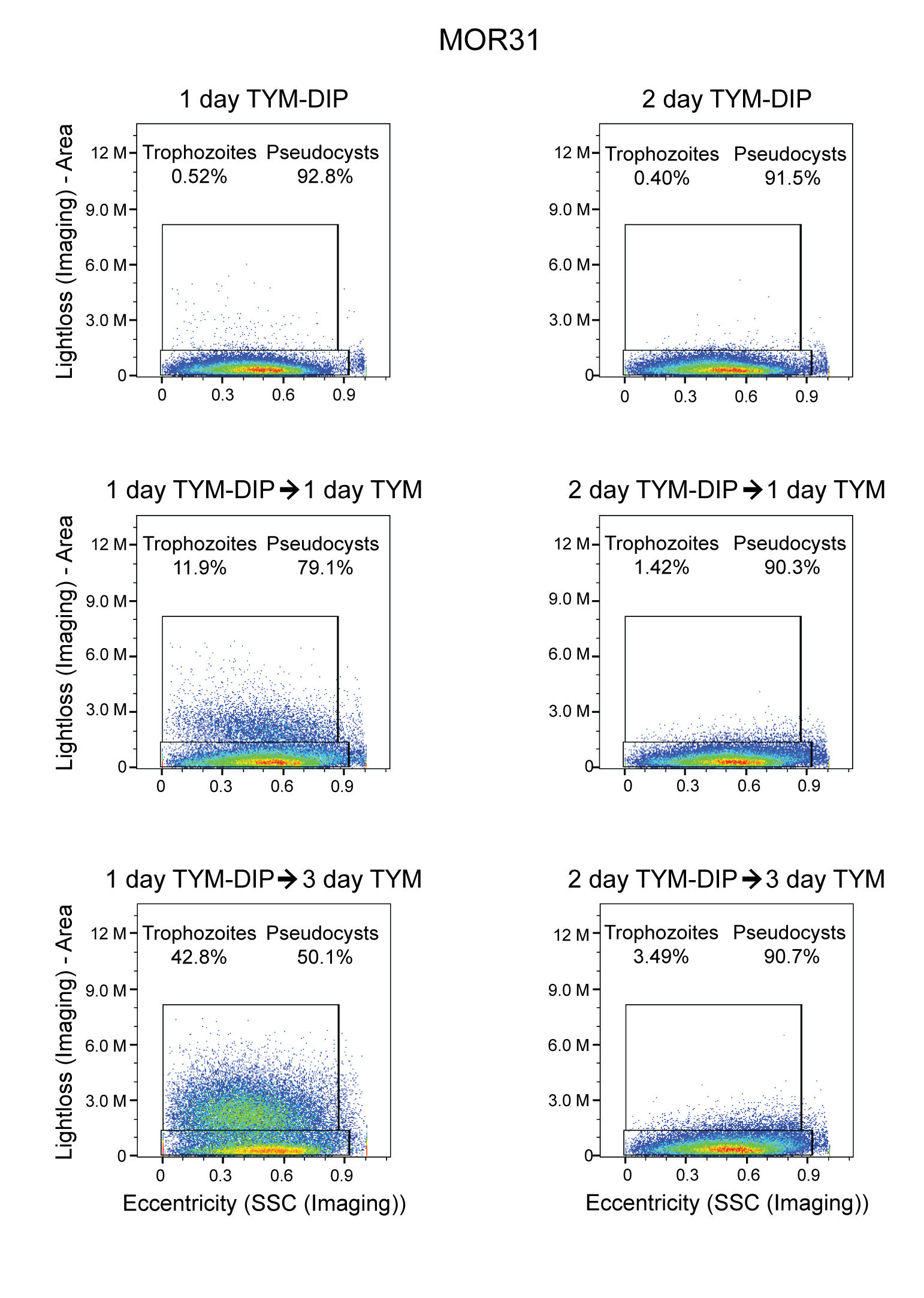

### Supplemental Fig 4

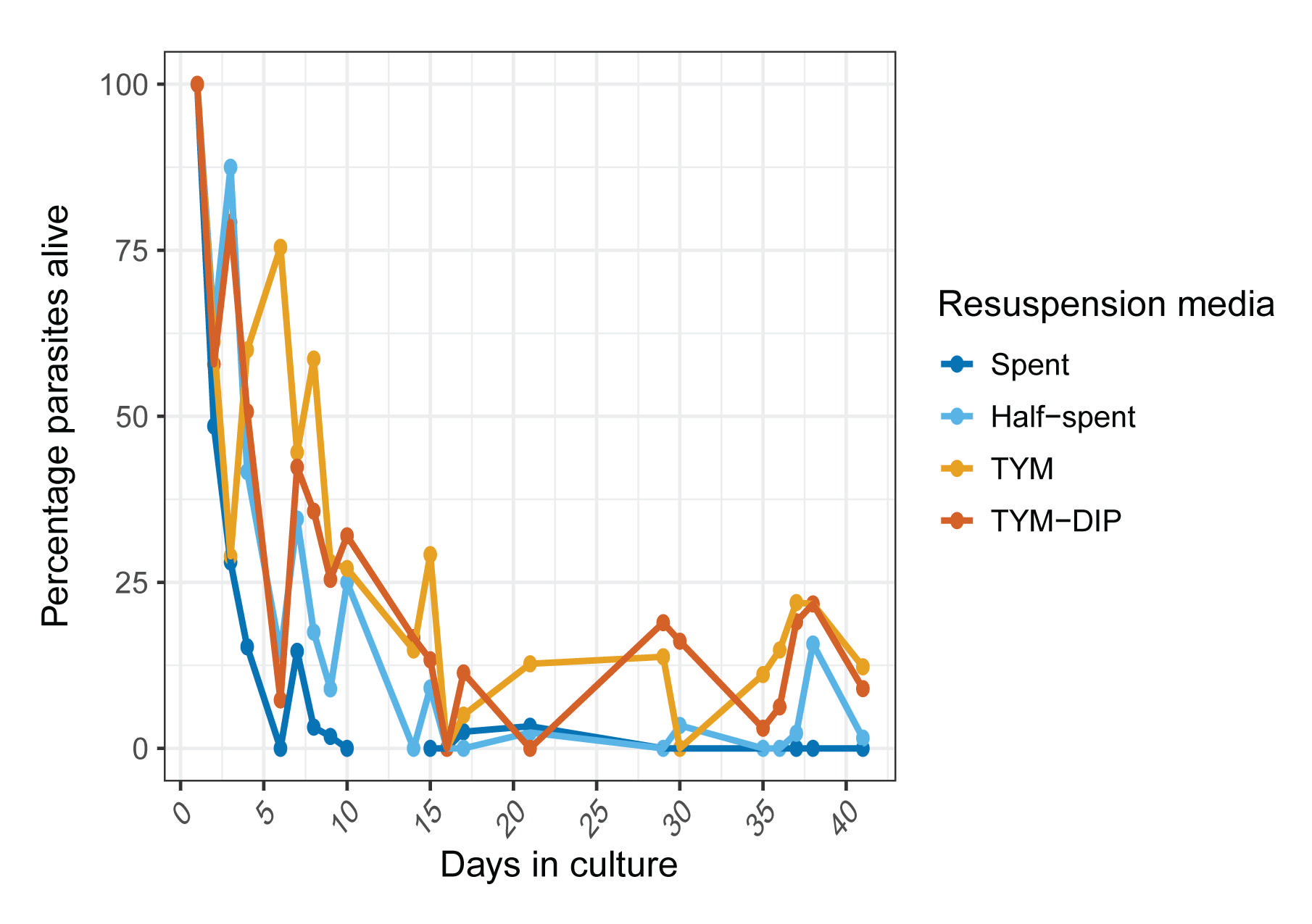

### Supplemental Fig 5

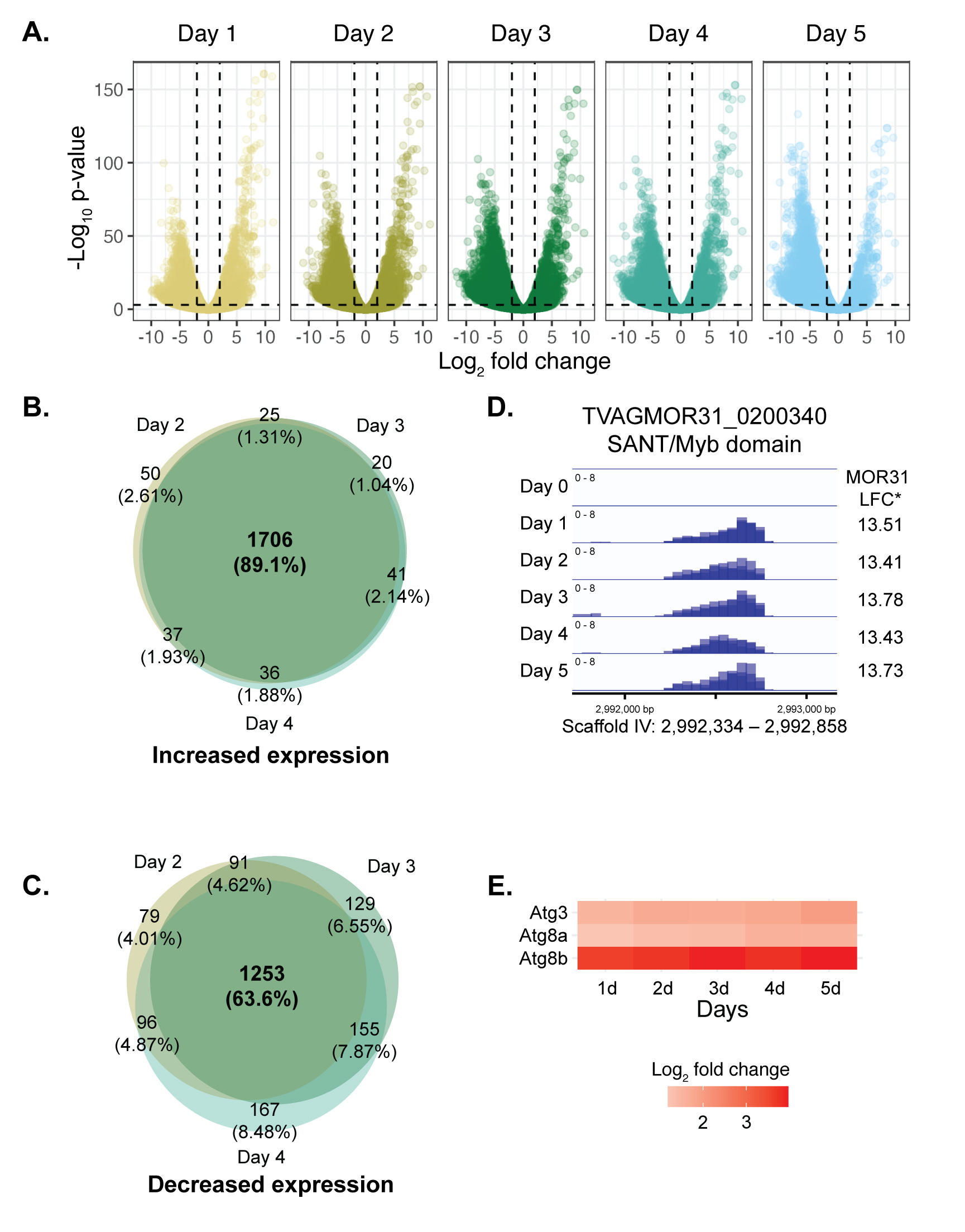
